# *In silico* analysis of piRNAs in retina reveals potential targets in intracellular transport and retinal degeneration

**DOI:** 10.1101/305144

**Authors:** Suganya Sivagurunathan, Nagesh Srikakulam, Jayamuruga Pandian Arunachalam, Gopal Pandi, Subbulakshmi Chidambaram

## Abstract

Long considered to be active only in germline, PIWI/piRNA pathway is now known to play significant role in somatic cells, especially neurons. Nonetheless, so far there is no evidence for the presence of piRNAs in the neurosensory retina. In this study, we have uncovered 102 piRNAs in human retina and retinal pigment epithelium (RPE) by analysing RNA-seq data. The identified piRNAs were enriched with three motifs predicted to be involved in rRNA processing and sensory perception. Further, expression of piRNAs in donor eyes were assessed by qRT-PCR. Loss of piRNAs in HIWI2 knockdown ARPE19 cells downregulated targets implicated in intracellular transport (SNAREs and *Rabs*), circadian clock (*TIMELESS*) and retinal degeneration (*LRPAP1* and *RPGRIP1*). Moreover, piRNAs were dysregulated under oxidative stress indicating their potential role in retinal pathology. Intriguingly, computational analysis revealed complete and partial seed sequence similarity between piR-62011 and sensory organ specific miR-183/96/182 cluster. Furthermore, the expression of retina enriched piR-62011 positively correlated with miR-182 in HIWI2 silenced Y79 cells. Thus, our data provides an evidence for the expression of piRNAs in human retina and RPE. Collectively, our work demonstrates that piRNAs dynamically regulate distinct molecular events in the maintenance of retinal homeostasis.

## Introduction

Retina, a neurosensory tissue of the eye, is developed from inner layer of optic cup regulated by eye field specific transcription factors (EFTF) whereas Retinal Pigment Epithelium (RPE) develops from outer layer of optic cup. Photoreceptors of the retina converts photons of light into electrical impulses where the crosstalk with RPE is necessary. Recent studies have indicated the involvement of non-coding RNAs in eye development plausibly by targeting the EFTF^1^. Besides their role in development, the importance of non-coding RNAs in visual function was confirmed when the depletion of miRNAs resulted in loss of outer segments of cone photoreceptors causing reduced light response^2^. Further, non-coding RNAs are also linked with retinal pathologies and are suggested to be novel targets for therapy^3^.

piRNAs are a group of small noncoding RNAs which bind with PIWI subclade of Argonaute family proteins^4–6^. They are 24 – 32 nucleotides (nt) long, methylated at their 3’ ends^7,8^ and are produced by a pathway independent of Dicer, making them distinct from other small noncoding RNAs. These group of RNAs are essential for maintaining the genomic integrity in germline cells by repressing the transposable elements^9^. Initially, it was thought to be restricted to germ cells, however, recent reports indicate their presence in various somatic tissues^10,11^ and cancer cells^12–14^. In addition, altered expression of piRNAs is observed during various pathological conditions like cardiac hypertrophy^15^ and liver regeneration^10^ in rats. Thus, growing evidence on the role of piRNAs in various non-germline tissues underscores the need to explore their importance in specific somatic cells as well.

Recent years has seen a surge of research findings on the existence of piRNAs in neurons. Initially, Lee *et al* has suggested roles for piRNAs in modulating the dendritic spine development in mouse hippocampal neurons^16^. In addition, altered expression of piRNAs were reported in rat brain after transient focal Ischemia^17^. Interestingly, a mechanistic role for piRNAs in neurons has been demonstrated by Rajasethupathy *et al* where PIWI/piRNA complexes in *Aplysia* enhanced memory-related synaptic plasticity through epigenetic regulation of CREB2^18^. Recently, piRNAs have shown to attenuate the axonal regeneration of adult sensory neurons in rats^19^. Moreover, piRNAs are deregulated in Alzheimer’s disease (AD) and are known to affect the disease associated pathways^20^. Nevertheless, there is no study on the presence and role of PIWI-interacting RNAs (piRNAs) in retina.

In our recent reports, we have shown that PIWI-like proteins are expressed in human retina and HIWI2 (PIWIL4) regulates tight junctions in maintaining the integrity of RPE^21^. HIWI2 also alters the expression of OTX2, an eye field specification marker in human retinoblastoma cells^22^. The presence of PIWI-like proteins in retina strongly implies the existence of piRNAs. Therefore, we investigated the expression and role of piRNAs in retina and RPE. The present study identified 102 piRNAs by computational analysis of RNA-seq data obtained from human retina and RPE tissues. Additionally, novel targets of piRNAs in multiple molecular processes have been validated. Our data presents new insights into the functions of piRNAs in retina and RPE.

## Results

### Expression of piRNAs in human retina and RPE

We examined whether piRNAs are expressed in retina and RPE by analysing the small RNA sequencing data retrieved from ArrayExpress (Accession ID: E-MTAB-3792)^23^. It consisted of 16 samples of human retina and 2 samples of RPE. The small RNA sequences were checked for their quality and the reads were mapped with the annotated small RNAs (tRNA, snRNA, snoRNA, miRNA, rRNA, mtRNA). The mapped reads were discarded to filter out the known small RNAs and the unmapped reads were aligned to the human genome. The resulting reads were considered as potential piRNA candidates and were mapped with the sequences of piRNAs downloaded from piRNAQuest^24^ (Fig. 1). Interestingly, we observed 102 piRNAs with tags per million (TPM) values > 1. piRNAs that were expressed in retina and RPE are listed in Table S1. A heatmap on the expression of the identified piRNAs indicated that piR-31068, piR-35982, piR-35284 were in abundance in both the tissues analysed. Interestingly, piR-62011, piR-36743 and piR-36742 were highly expressed only in retina. Similarly, the expression of piR-35411, piR-35412, piR-35413, piR-31104 and piR-31103 were higher only in RPE tissue (Fig. 2a & b). Together, the results showed that piRNAs were expressed in ocular tissues and indeed differential expression of piRNAs were observed between retina and RPE.

**Fig 1.**
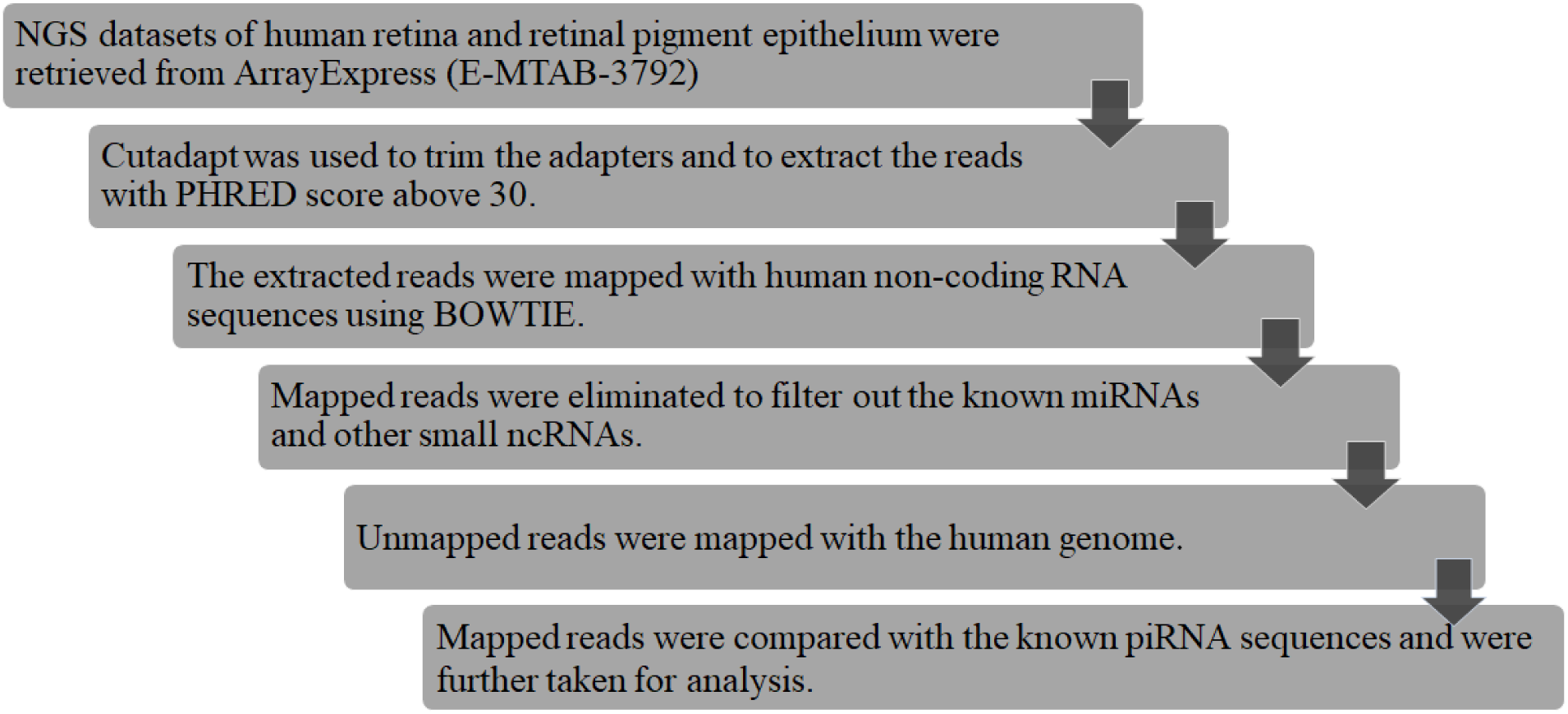
Workflow of piRNA identification. The flowchart describes the step by step protocol of piRNA analysis using the small RNA sequencing dataset of human retina and retinal pigment epithelium.

**Fig 2.**
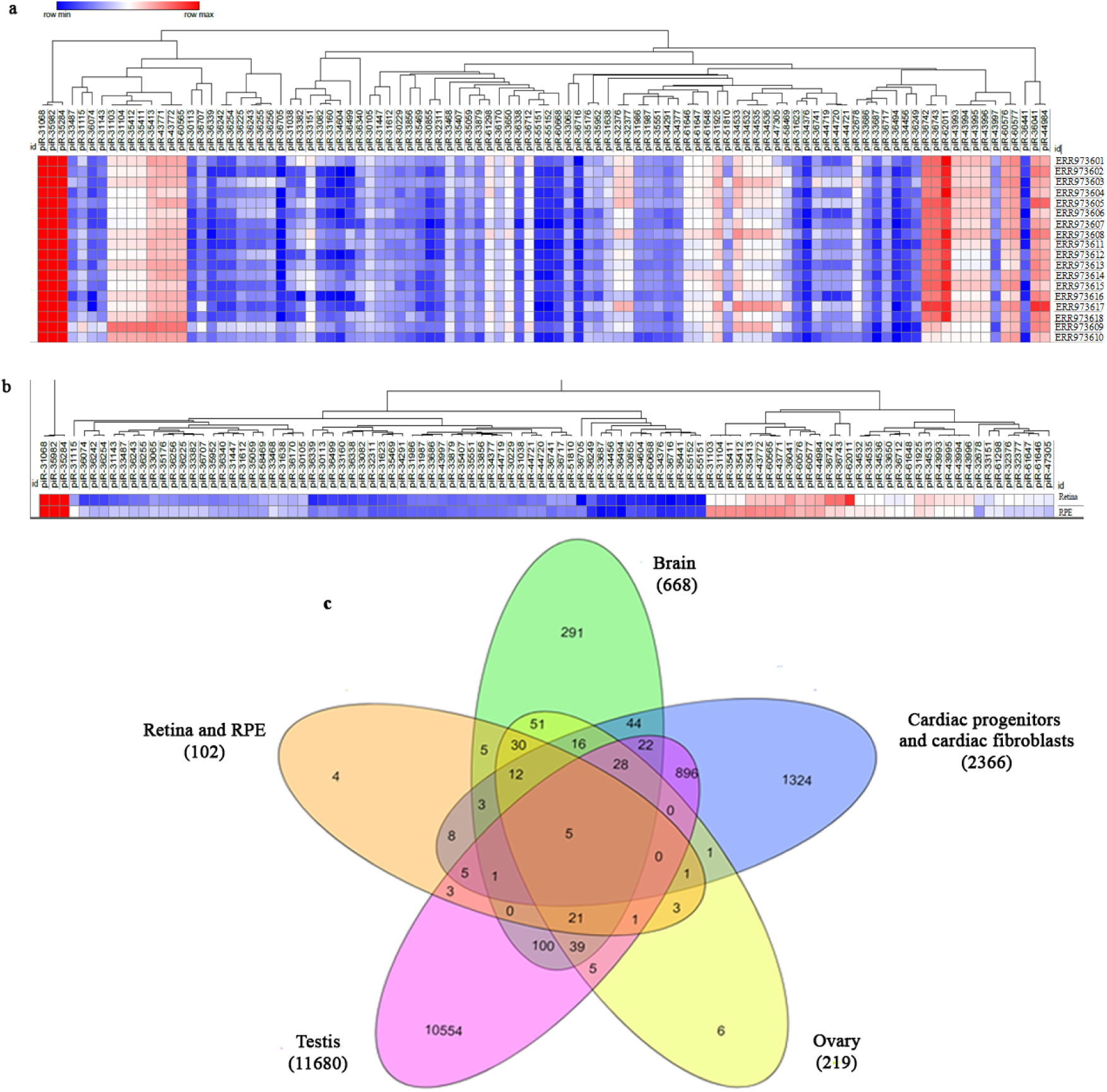
In silico analysis for identification of piRNAs in human retina and retinal pigment epithelium (RPE). a) Heatmap shows the expression of the identified 102 piRNAs in human retina (16 datasets) and RPE (2 dataset). b) Heatmap depicts the differential expression of piRNAs in retina and RPE. Tags per million (TPM) values are used to generate heatmap using Morpheus. c) Venn diagram depicts the unique and commonly expressed piRNAs in 6 human tissues, retina and RPE; brain; cardiac progenitors and cardiac fibroblasts; ovary and testis.

### Tissue enriched piRNAs

Having observed the piRNAs in ocular tissues, we next investigated whether the identified piRNAs are tissue specific. To this end, our data was compared with the piRNAs so far reported in human brain, cardiac progenitors and cardiac fibroblasts, ovary and testis^20,25–27^. We observed 4 piRNAs (piR-30855, piR-34535, piR-44719 and piR-61647) to be expressed only in retina and RPE. All the analysed tissues shared 5 piRNAs (piR-33382, piR-33468, piR-35284, piR-35413 and piR-36225) in common. Moreover, 3 piRNAs (piR-32376, piR-43771 and piR-62011) were found to be commonly present in retina, RPE and testis. However, piR-62011 is abundantly expressed in retina (TPM value - 112874) when RPE is considered (TPM value - 1003) (Fig 2b). Notably, around 75% of the piRNAs in the retina/RPE tissues matched with the piRNAs in brain, followed by 72% with ovary. In contrast, piRNAs of ocular tissues showed only 35% and 34% match with testis and cardiomyocytes (cardiac progenitors and cardiac fibroblasts) respectively (Fig. 2c). Thus, a significant overlap was found amongst piRNAs expressed in human brain and retina/RPE; and piR-62011 appeared to be a retina enriched piRNA.

### Genomic origin of piRNAs

piRNAs are shown to originate originally from the transposon sequences. The origin of the identified piRNAs indicated that majority of them mapped to tRNA family repeats, especially to tRNAs Val and Gly followed by Ala. This is in accordance with the earlier report on somatic piRNAs derived from the tRNA fragments^28^. Moreover, protein coding genes were also known to be a source of piRNAs^29^ and likewise our data specified 28 protein coding genes as origin for a set of piRNAs. Besides, 20 of the piRNAs originated from SNORD gene family which encodes small nucleolar RNAs (snoRNA). Recently, small nucleolar RNAs were reported to be precursor of piRNA^30^. Strikingly, the retina enriched piR-62011 had its origin in the 3’ UTR of miR-182 (Fig. 3a & b, Table S2).

**Fig 3.**
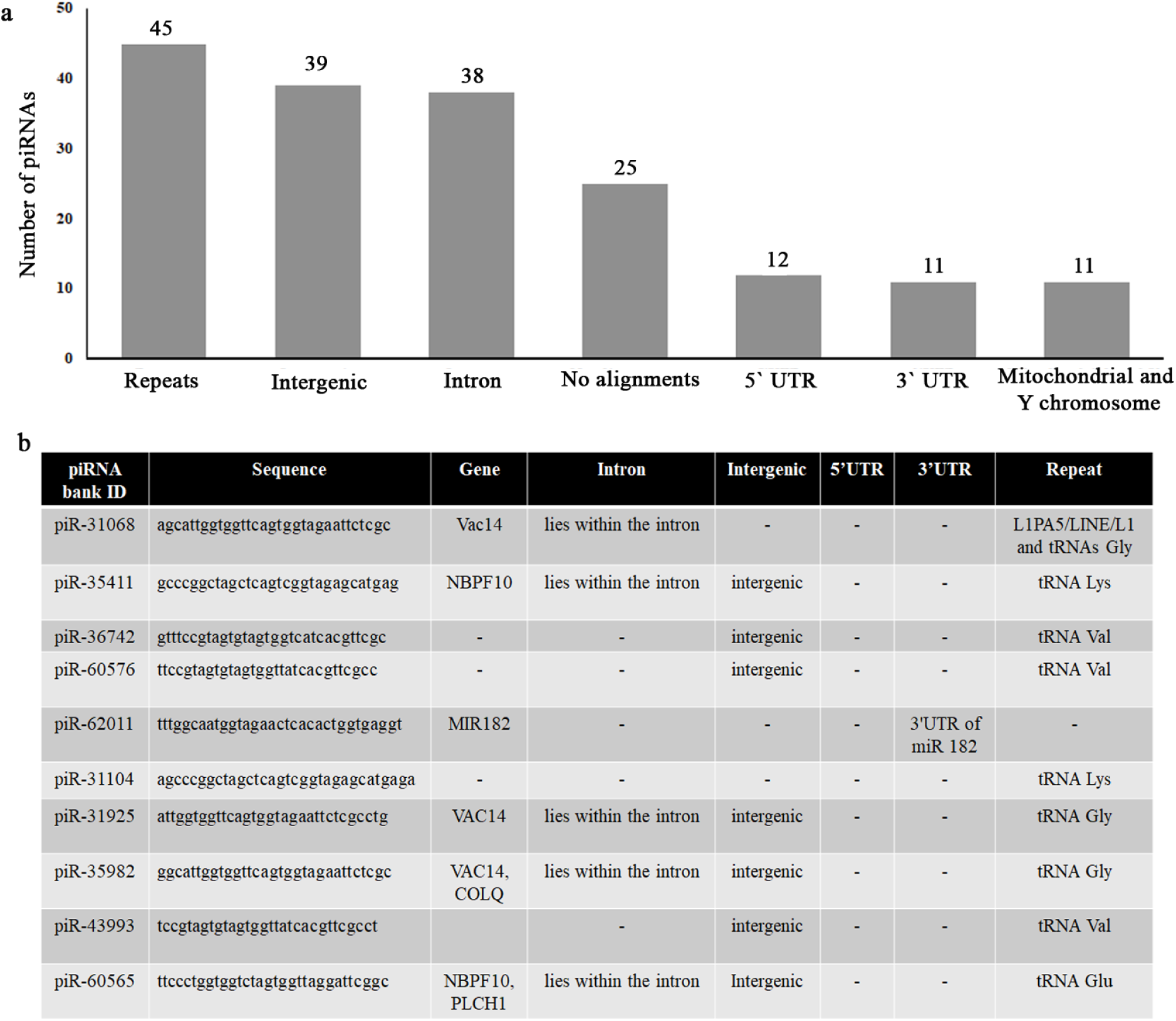
Genomic origin of piRNAs. a) Bar graph depicts the data obtained from piRNAQuest with the number of piRNAs that originate from the mentioned genomic regions. b) A representative list of identified piRNAs with piRNABank ID, sequence and genomic origin are presented in the table. This list also includes the 5 piRNAs validated in the study.

Unlike the piRNAs expressed in germline cells, somatic piRNAs are reported to map ‘piRNA clusters’ to a lesser extent^31^. In order to understand the genomic localization of piRNAs in retina/RPE, we considered minimum 3 piRNAs originating within 20 kb distance to form a cluster. Our analysis indicated that chromosome (chr) 6 harboured four piRNA clusters followed by chr 1 with three clusters. Chr 2, 5 and 16 had two piRNA clusters in each while chr 3,11,14,17 and 18 expressed one piRNA cluster. Interestingly, 2 piRNA clusters had its origin in mitochondrial genome, which is in line with the recent identification of piR-36707 and piR-36741 in mitochondria of the mammalian cancer cells^32^ (Table 1).

**Table 1.** Chromosomal distribution of piRNAs.

| Chromosome | Number of piRNA clusters | Number of piRNAs in each cluster |
| --- | --- | --- |
| **Chromosome 1** | 3 | Cluster 1 – 6 piRNAs<br>Cluster 2 – 4 piRNAs<br>Cluster 3 – 5 piRNAs |
| **Chromosome 2** | 2 | Cluster 1 – 5 piRNAs<br>Cluster 2 – 4 piRNAs |
| **Chromosome 3** | 1 | Cluster 1 – 3 piRNAs |
| **Chromosome 5** | 2 | Cluster 1 – 5 piRNAs<br>Cluster 2 – 9 piRNAs |
| **Chromosome 6** | 4 | Cluster 1 – 7 piRNAs<br>Cluster 2 – 7 piRNAs<br>Cluster 3 – 5 piRNAs<br>Cluster 4 – 4 piRNAs |
| **Chromosome 11** | 1 | Cluster 1 – 4 piRNAs |
| **Chromosome 14** | 1 | Cluster 1 – 3 piRNAs |
| **Chromosome 16** | 2 | Cluster 1 – 6 piRNAs<br>Cluster 2 – 4 piRNAs |
| **Chromosome 17** | 1 | Cluster 1 – 5 piRNAs |
| **Chromosome 18** | 1 | Cluster 1 – 3 piRNAs |
| **Mitochondrial genome** | 2 | Cluster 1 – 6 piRNAs<br>Cluster 2- 4 piRNAs |
| **Chromosome 4, 7, 8, 9, 10, 12, 13, 15, 19,20, 21, 22, X and Y chromosome** | No clusters. | 0-4 piRNAs |

### Enriched motifs

Though piRNAs are not conserved across species, these small RNAs have been observed to possess short motifs. We identified 3 conserved motifs in piRNAs using motif predictor MEME and the first motif (21 nt long) with an E-value of 5.4 e^-127^ was conserved in 38 piRNA sequences. Further, the function and localization of the nucleotide motif was predicted using GOMo which showed that the nucleotide sequences in the first motif was found to be involved in rRNA processing and localized to nucleolus and cytoplasm. The second motif (21 nt long) with an E-value of 1.3 e^-023^ was conserved in 14 piRNAs and was predicted to be localized in mitochondria. The third motif (28 nt long) with an E-value of 2.2 e^-016^, speculated to reside in extracellular region, was conserved in 21 piRNA sequences. This particular motif is predicted to be involved in G-protein coupled protein receptor signalling pathway and sensory perception (Fig. 4a).

**Fig 4.**
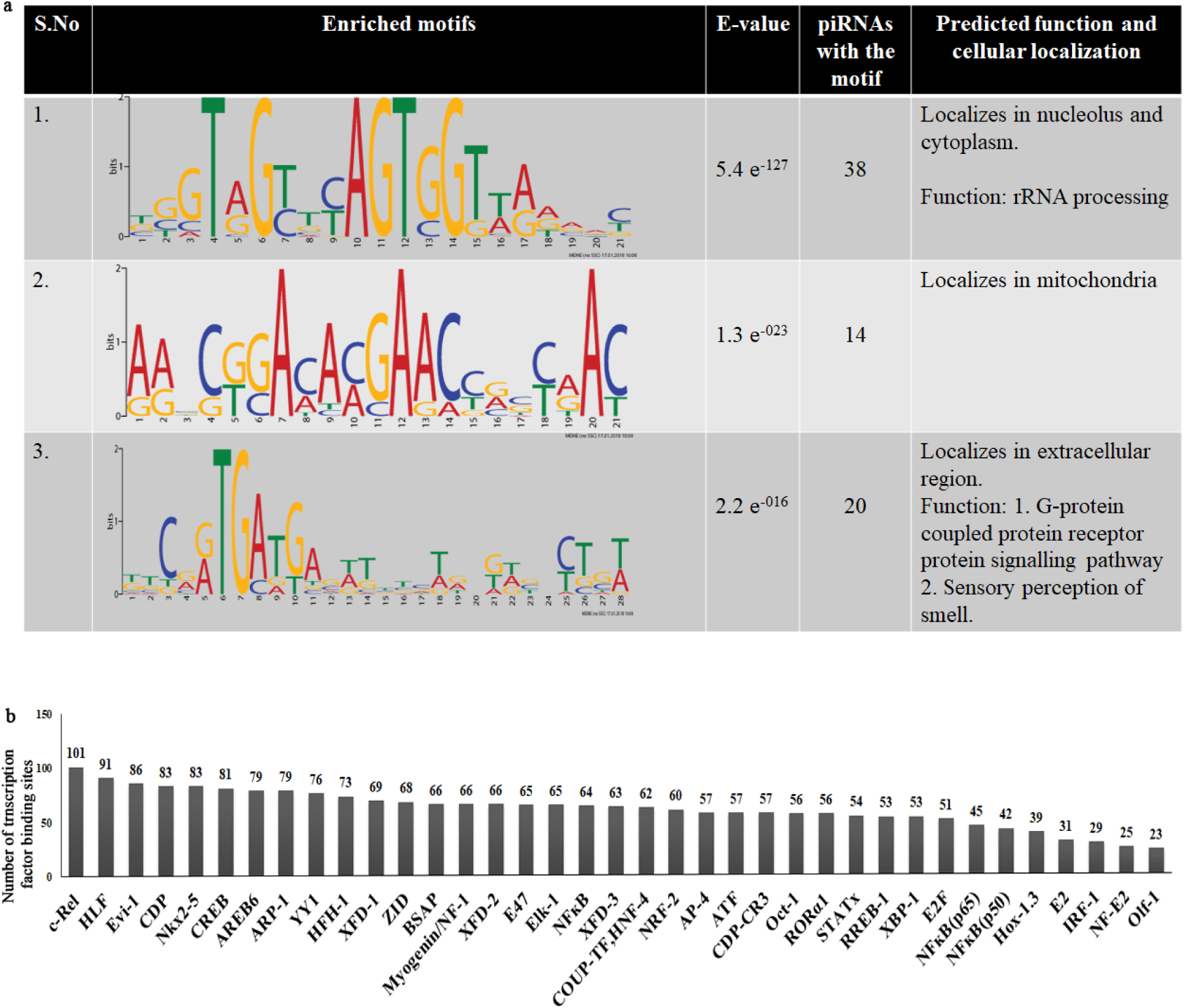
Enriched motifs and regulators of piRNAs. a) Logos with the corresponding E-values represent the motifs identified by MEME. Function and cellular localization of the motifs predicted by GOMo are also shown. b) Bar graph represents the number of transcription factor binding sites observed in the upstream sequences of 102 piRNAs.

### Transcription factors regulating piRNAs

Given the presence of piRNAs in human retina and RPE, we further attempted to describe the possible regulatory factors of piRNA. Sequences upstream of the piRNA loci were searched for the presence of transcription factor binding sites (Fig. 4b). It is interesting to note that transcription factor c-Rel showed higher number of binding sites in the promoters of piRNAs followed by Hepatic Leukemia factor (HLF). c-Rel is a member of NFĸB transcription factor family and we also observed binding sites for NFĸB in the promoters of piRNA. Though c-Rel defects are mainly associated with inflammatory disorders they are also associated with memory and synaptic plasticity^33^. Meanwhile, piRNAs were also reported to be crucial for synaptic plasticity and memory storage by regulating CREB2^18^. Our data indicated that there were binding sites for CREB in the promoter regions of the piRNAs. c-Rel, HLF, Evi-1, CCAAT-displacement protein (CDP) and Nkx2-5 were the transcription factors which contained the most number of binding sites in the analysed regions.

### Evaluation of the identified piRNAs in human retina and RPE samples

Out of the 102 identified piRNAs, we evaluated the expression of piR-31068 (abundant in both retina and RPE), piR-35411 (abundant in RPE), piR-36742, piR-62011(abundant in retina) and piR-60576 (moderate expression in both the tissues). qPCR was done in human retina and RPE tissues dissected from donor eyes which confirmed the expression of these piRNAs (Fig. 5a). cDNA conversion without the addition of reverse transcriptase (or) the cDNA that were converted using random primers did not result in amplification indicating the specificity of the reaction. Among the piRNAs that were evaluated, piR-31068 is abundantly expressed (with low ΔCq), which is in accordance with the TPM values obtained (Table S1).

**Fig 5.**
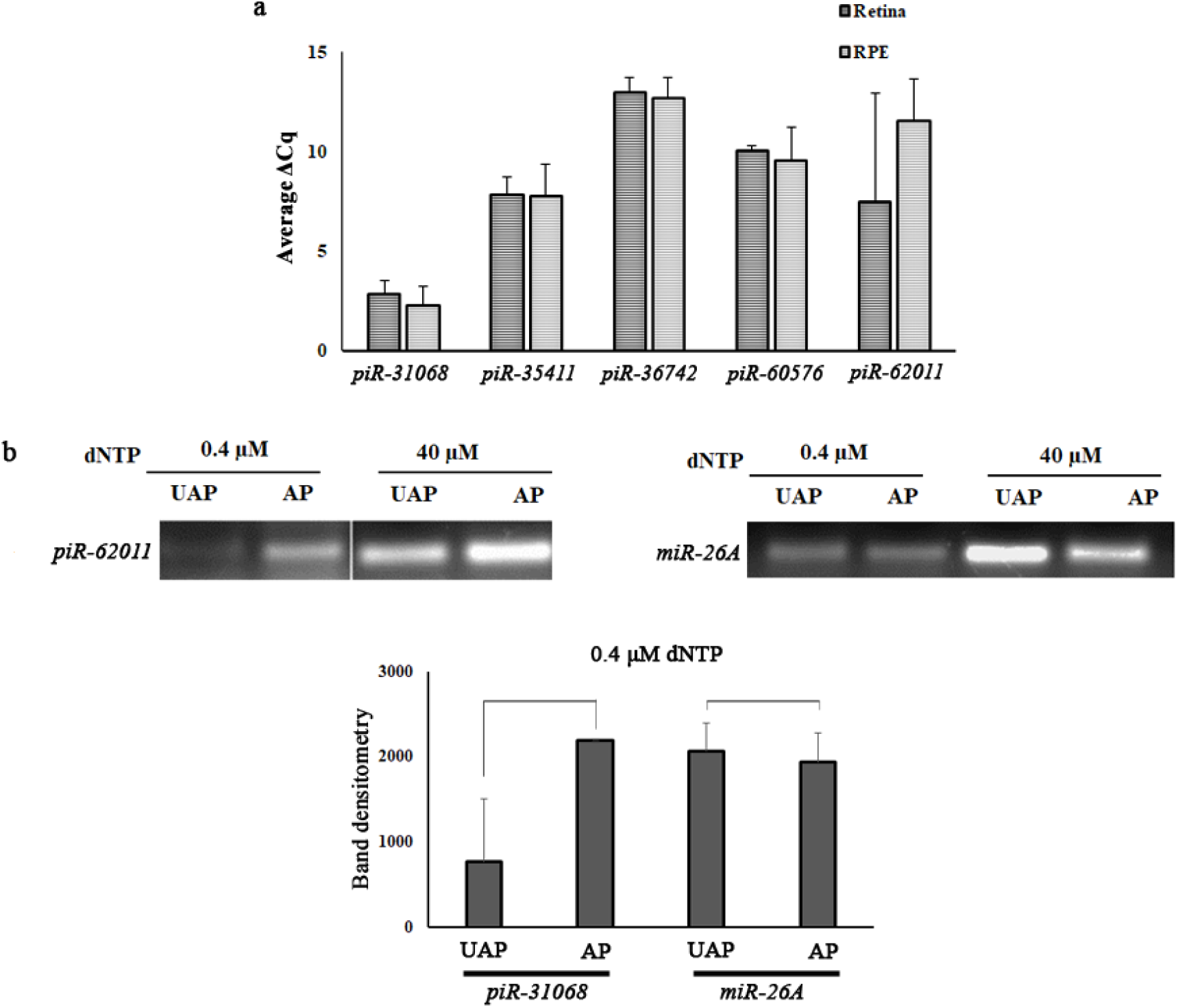
Validation of piRNA expression in human retina and RPE tissues. a) Bar graph represents the average ΔCq values of piRNAs (*piR-31068, piR-35411, piR-36742, piR-60576 and piR-62011*) obtained using qRT-PCR from retina and RPE tissues. Since the bar graph shows the average ΔCq values, the smaller bar represents higher expression and vice versa. The Cq values were normalized with U6. b) RTL-P analysis confirms the 3’-terminal 2’-O-methylation of piRNAs. Amplification of piRNA with unanchored primer at low dNTP concentration was reduced whereas the miRNA did not show any significant difference between unanchored and anchored primers. Full length images of the gels are presented in supplementary figure S1. (RTL-P – Reverse Transcription at Low dNTP concentration followed by PCR; UAP – Unanchored primer; AP – Anchored primer)

To validate whether the amplicons are indeed piRNAs, we used Reverse Transcription at Low dNTP concentration followed by PCR (RTL-P) analysis. This method investigated 2’ O - methylation at the 3’ ends of piRNAs. Reverse transcriptase stalls at methylated sites at low dNTP concentration. Exploiting this feature of the enzyme, piRNAs and miR-26a (control) were amplified at both high and low concentration of dNTP. During amplification, two different primers were used – unanchored^34^ (3’ methylation site of the piRNA will not be covered) and anchored (3’ methylation site of the piRNA will be covered by the primer) primers. Amplification of piRNAs in the presence of unanchored primer at low dNTP concentration was low whereas it was more at high dNTP concentration. miR-26a which was not methylated at its 3’ ends did not show such changes (Fig. 5b). Hence, the methylation at 3’ ends of the amplicons further confirmed that the obtained bands were indeed piRNAs. Altogether, our data confirmed the expression of piRNAs in human retina and RPE tissues.

### Putative targets of piRNAs

We used miRanda algorithm to identify the putative targets of piRNAs, to gain insight into the possible regulatory function. We obtained a list of potential targets with a threshold score of 200 and a minimum energy value of −20 kCal/Mol (Table S3). When the targets were functionally annotated, we observed that 24% of them were uncharacterised proteins. Majority of the putative targets were channels, transporters and proteins necessary for signal transduction followed by 13% and 14% of proteins essential for metabolism and nucleotide binding respectively. Around 9% of the proteins are involved in intracellular trafficking and 7% belongs to proteins important for adhesion, cytoskeletal arrangements and cell-cell contact including tight junction proteins. The other predicted targets are known to be crucial for immune and stress response, sensory perception, neuronal activity and DNA repair (Fig. 6a).

**Fig 6.**
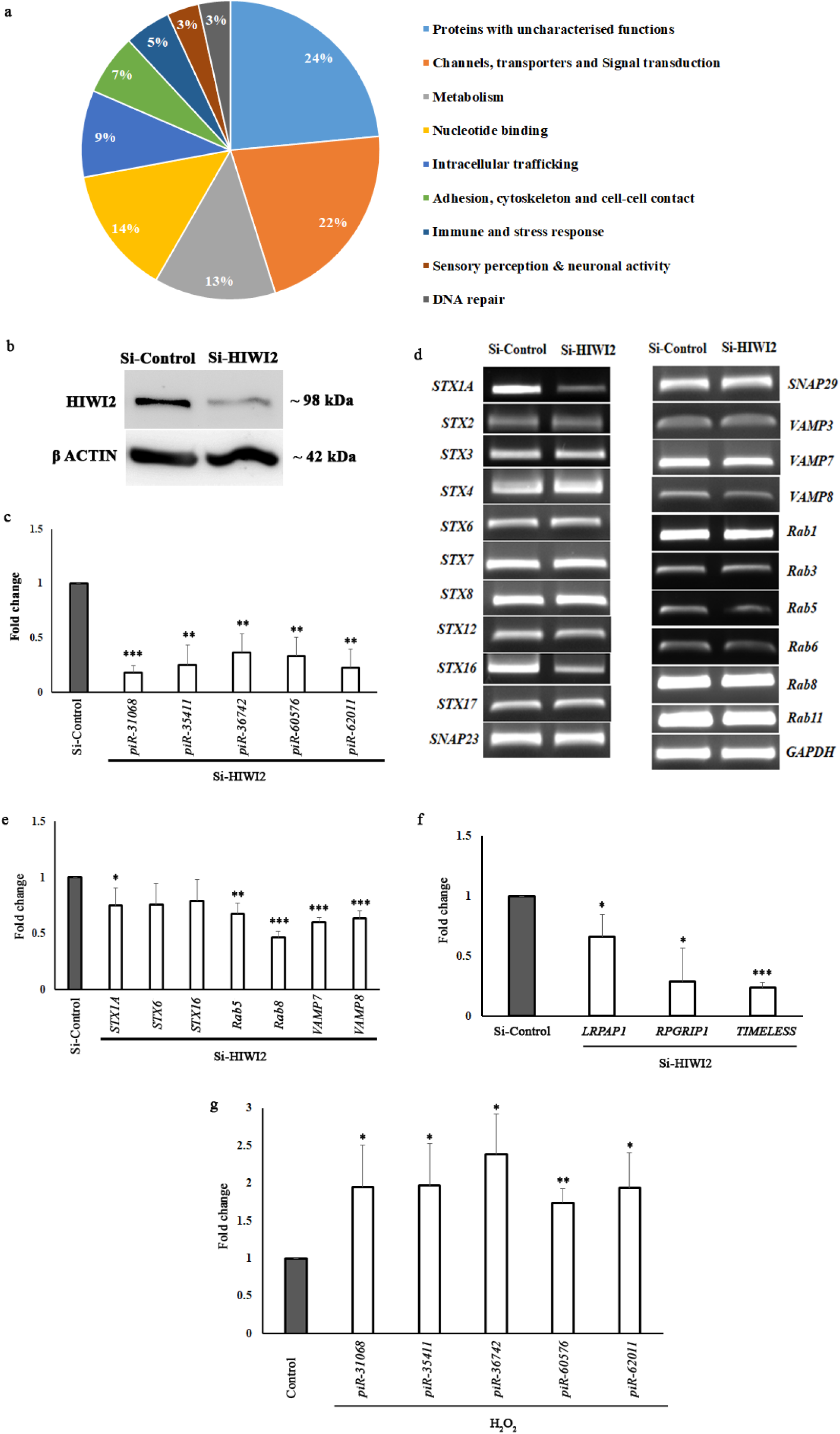
Potential targets of piRNAs. a) Pie chart shows the percentage of putative targets of piRNA in each category predicted by miRanda algorithm. b) Downregulation of HIWI2 is shown by Western blot in Si-Control and Si-HIWI2 ARPE19 cells. βACTIN is used as loading control. Full length images of the blots are presented in supplementary figure S2. c) Expression of piRNAs in Si-HIWI2 ARPE19 cells is shown as fold change obtained by qRT-PCR. U6 levels are used for normalization. d) Agarose gel image showing the expression of *SNARE*s and *Rab*s in Si-Control and Si-HIWI2 cells. *GAPDH* is used as the loading control. e) Bar graph represents the fold change of *STX1A, STX6, STX16, Rab5, Rab8, VAMP7* and *VAMP8* in Si-HIWI2 ARPE19 cells obtained by qRT-PCR. *18S rRNA* is used for normalization. f) Bar graph shows the fold change of *LRPAP1, RPGRIP1* and *TIMELESS* in Si-HIWI2 ARPE19 cells. The expression is normalized with *18S rRNA*. g) Fold change of piRNAs obtained by qRT-PCR from the cells subjected to oxidative stress using 200 µM H_2_O_2_ are depicted as bar graph. U6 is used for normalization. (\**p*<0.05; \*\**p*<0.01; \*\*\**p*<0.001; Student’s *t* test was used to analyse the statistical significance between Si-Control and Si-HIWI2; control and H_2_O_2_ treated cells)

### Evaluation of the predicted targets of piRNA

Among the predicted targets, the possible influence of piRNAs in cell signalling and cell adhesion proteins can be conjectured from our previous study^21^. We demonstrated the function of HIWI2 in altering the tight junction proteins through Akt/GSK3β signalling pathway^21^. Since PIWI proteins bind with piRNAs, effect of HIWI2 on tight junctions might be plausibly through piRNAs. Next, we analysed whether absence of piRNAs influenced the expression of putative targets. HIWI2, a ubiquitously present member of 4 PIWI-like proteins in human, was efficiently silenced in ARPE19 (RPE) cell line as indicated by the Western blot (Fig. 6b). Accordingly, the piRNAs were downregulated in Si-HIWI2 cells (Fig. 6c) and the interesting novel targets from intracellular trafficking machinery were analysed. We screened for the expression of *SNARE*s and *Rab* transcripts upon HIWI2 silencing in ARPE19 by semi RT-PCR (Fig. 6d). Further qRT-PCR indicated that *STX1A*, *Rab5, Rab8, VAMP7* and *VAMP8* were significantly downregulated in Si-HIWI2 cells (Fig. 6e). In addition, the expression of other predicted targets such as *TIMELESS*, *RPGRIP1* and *LRPAP1* were also evaluated. Strikingly, *TIMELESS* transcripts were 4.34-fold downregulated in the absence of HIWI2 (Fig. 6f). Both, *RPGRIP1* and *LRPAP1* transcripts were 3.57-fold and 1.51-fold downregulated in Si-HIWI2 cells respectively (Fig. 6f). Notably, it was reported that *RPGRIP1* mutant of Zebrafish did not develop rod outer segments in the retina and were suggested to regulate ciliary protein trafficking^35^. *LRPAP1* is known to be associated with myopia^36^ and the variations in LRPAP1 were observed to be associated with late onset of Alzheimer’s disease^37^. Thus, loss of piRNAs downregulated the expression of *SNARE*s, *Rab*s and *RPGRIP1* which are important for ciliary trafficking. In addition, *LRPAP1* and the expression of key circadian clock protein, *TIMELESS* were also reduced in the absence of piRNAs.

### Elevated levels of piRNA under oxidative stress

Oxidative stress is a major cause in the pathogenesis of ocular diseases such as keratoconus, cataractogenesis and age-related macular degeneration^38^. To gain insight into the importance of piRNAs in pathological conditions, ARPE19 cells were subjected to oxidative stress. Upon treatment of the cells with 200 μM of H_2_O_2_, the piRNAs were significantly upregulated. We observed 2.3-fold upregulation of piR-36742 followed by 1.9-fold increase in piR-35411, piR-31068, piR-62011 and 1.7-fold upregulation in piR-60576 (Fig. 6g). Clearly, the expression of piRNAs were altered under oxidative stress.

### Alignment of piRNA with miRNA and long non-coding RNA (lncRNA)

Since the crosstalk between the non-coding RNAs is evolving^39^, we investigated the miRNAs and lncRNAs that might be related to the identified piRNAs. Alignment of the piRNA sequences with miRNAs showed that piR-35059, piR-36741 and piR-62011 matched with the seed sequences of miR-4420, miR-4284 and miR-182 respectively (Fig 7a). It was interesting to note that piR-62011, an abundant piRNA in retina showed complete sequence homology to miR-182, which was reported to be the highly expressed miRNA in human retina^23^. Further, to understand whether the expression of piR-62011 and miR-182 are related, we examined whether silencing HIWI2 (thereby the loss of piRNAs) alters the expression of miR-182 in Y79, a retinoblastoma cell line. Remarkably, the expression of both piR-62011 and miR-182 were reduced 1.30 and 3.13- fold in HIWI2 silenced Y79 cells respectively (Fig 7b & c).

**Fig 7.**
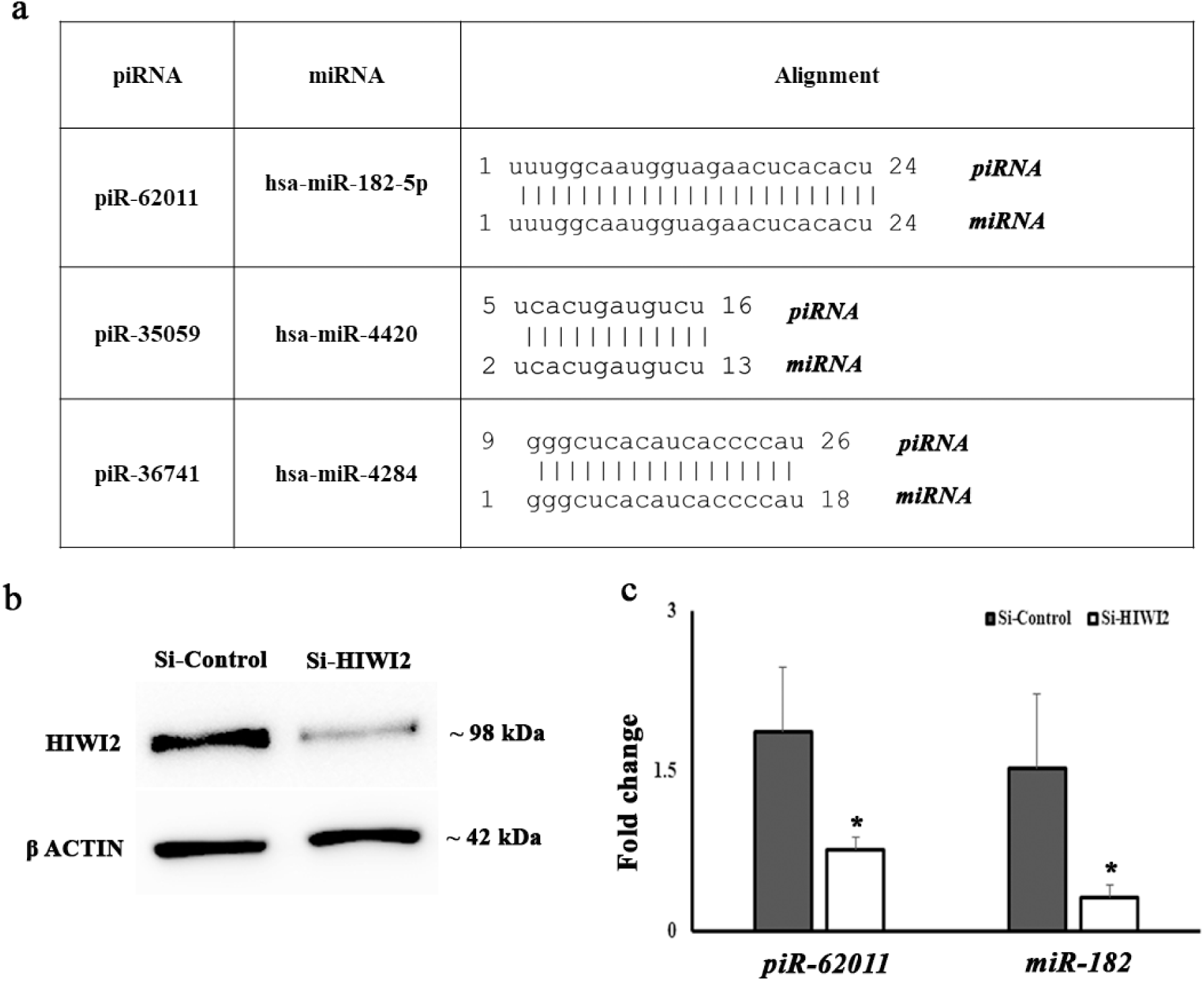
miR-182 is downregulated in Si-HIWI2 Y79 cells. a) The table lists the miRNAs which showed seed sequence similarity with piRNAs. b) Western blot shows the expression of HIWI2 in Si-Control and Si-HIWI2 Y79 cells. βACTIN is used as loading control. Full length images of the blots are presented in supplementary figure S2. c) Bar graph represents the fold change of *piR-62011* and *miR-182* obtained by qRT-PCR in Si-Control and Si-HIWI2 Y79 cells. U6 is used for normalization. (\**p*<0.05; Student’s *t* test was used to analyse the statistical significance between Si-Control and Si-HIWI2)

**Fig 8.**
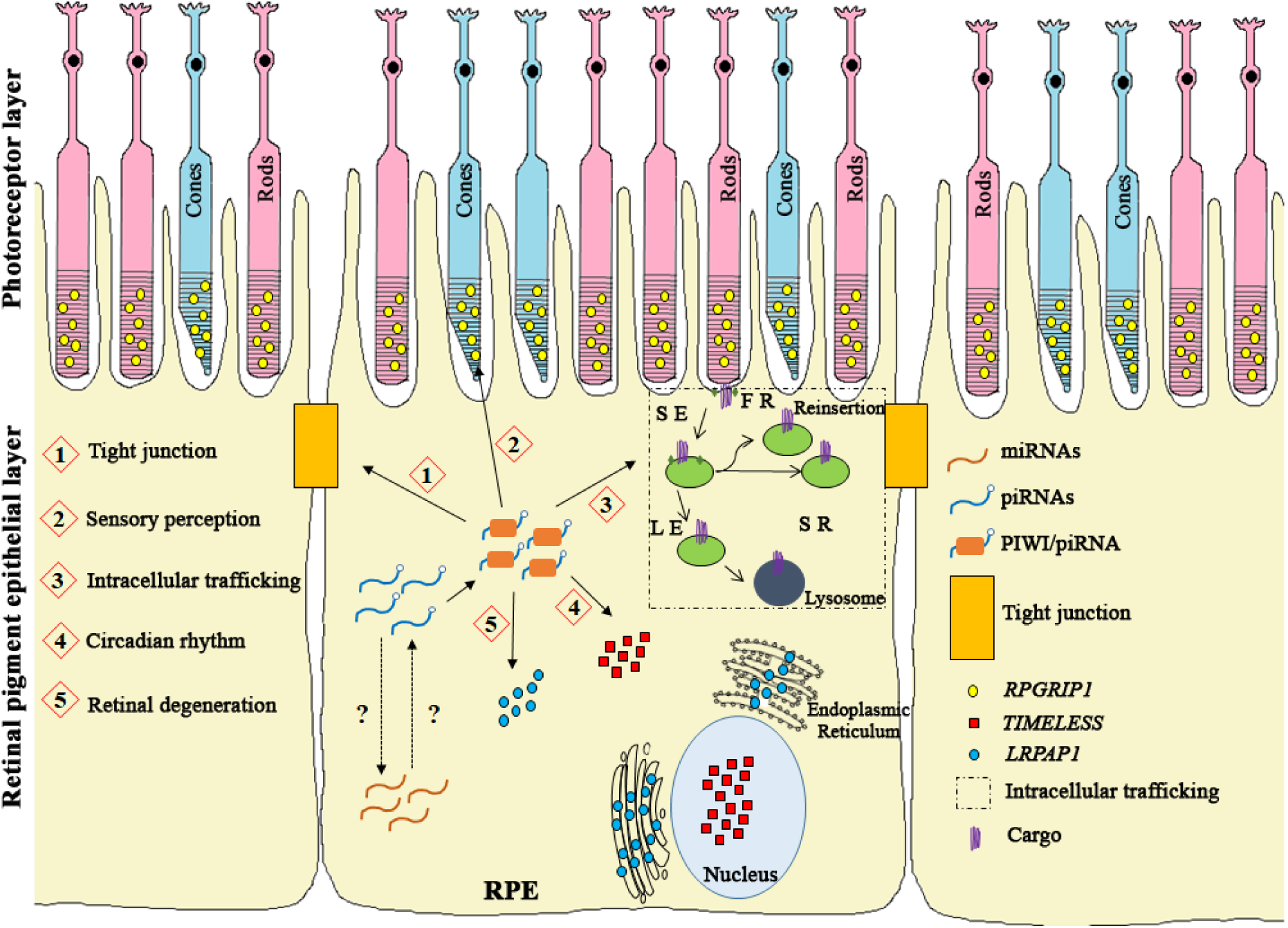
Hypothetical model for pleiotropic functions of piRNAs. The model depicts the multiple functions of piRNAs in RPE in (1) Altering the expression of tight junctions; (2) Regulation of ciliary trafficking and sensory perception by influencing *RPGRIP1;* (3) Intracellular trafficking by affecting the expression of *STX1A, Rab5, Rab8, VAMP7 and VAMP8*; (4) Regulation of circadian clock by acting on *TIMELESS*; (5) Involvement in retinal degeneration through modulation of *LRPAP1* and *RPGRIP1*. Regulation of these processes by piRNAs may include a functional interaction with miRNAs. Gaining deeper understanding of the exact molecular mechanism will expand our knowledge on pleiotropic effects of piRNAs in somatic cells. (SE: Sorting Endosome; FR: Fast Recycling; SR: Slow Recycling; LE: Late Endosome)

Additionally, some piRNAs showed an exact sequence match with 20 lncRNAs and few with either 1 or 2 mismatches. Many of the lncRNAs that mapped with the piRNAs were uncharacterised (Table S4). Apparently, the sequence similarities between miR/piR/lncRNA hint a potential crosstalk among the non-coding RNAs in the retina.

## Discussion

Recently, we reported the presence of PIWI-like proteins in ocular tissues and the importance of HIWI2 in the maintenance of epithelial integrity of RPE. Since PIWI proteins are associated with piRNAs, it was compelling to examine whether piRNAs play a role in retinal function. The present study confirmed the expression of piRNAs in retina and RPE. Moreover, loss of piRNAs altered the expression of specific transcripts involved in intracellular trafficking, sensory perception and circadian rhythm. We have also observed elevated levels of piRNAs under oxidative stress denoting their association with pathological conditions.

Our analysis showed that majority of the piRNAs in retina and RPE originate from tRNA fragments of Val and Gly, which is in line with the findings in human brain^20^ and ovarian cancer tissues^26^. Although, recent studies demonstrated the existence of small nucleolar-RNA (sno-RNA) derived piRNAs and a novel pathway for gene regulation^30,40^, still, far less is known about this group of piRNAs. The SNORD family derived piRNAs in retina/RPE may play important functions which warrants further investigation.

It was surprising to find that piR-62011, an abundant piRNA in retina displayed seed sequence match and aligned completely with miR-182. Notably, seed sequence similarity between miR-17-5p and piRNAs has been shown to reciprocally regulate each other during mouse embryonic development^41^. However, the expression of miR-182 positively correlated with piR-62011. Rinck *et al* has shown that binding of multiple miRNAs with a single mRNA may either have independent or cooperative regulation among them^42^. Likewise, in a recent study piRNAs have been shown to act similar to miRNAs in targeting the mRNA with cooperative regulation^43^. Taken together, based on the expression of piR-62011 and miR-182, they may have cooperative activity rather than reciprocal regulation. This can be further justified by the fact that the binding sites of piR-62011 and miR-182 predicted by RNA Hybrid^44^ in few selected target mRNAs, varied from very distal to very close or adjacent implying either independent or cooperative regulation between piRNAs and miRNAs. The presence of two classes of small non-coding RNAs with similar sequences points out a possibility that PIWI could bind and facilitate the miRNAs. This notion can be substantiated by the recent finding that PIWI is required for the expression of a subset of miRNAs in mouse ^45^.

Notably, miR-183/96/182 is reported as ‘sensory organ specific’ miRNA cluster abundantly expressed in mouse retina^46^. Moreover, this cluster is modulated by light-dark adaptation^47^. In support of these findings, absence of miR-182 and miR-183 lead to the loss of cone outer segments and were also described to be necessary for the formation of inner segments, connecting cilia and short outer segments of photoreceptor in adult mice^2^. Moreover, miR-183/96/182 has been downregulated in a mouse model for inherited blindness^48^. Further, miR-182 is associated with an ocular pathology called primary open-angle glaucoma^49^. Hence, we speculate an independent or co-operative regulation between piR-62011 and the sensory organ specific miR-183/96/182 through seed sequence match. It would be relevant to investigate the role of piR-62011 in glaucoma and retinal degeneration.

It is interesting to note the three conserved motifs in retina and RPE. Earlier, Grivna *et al* have observed an 8-nt motif shared by 17 sequences and a 21-nt motif shared by 5 sequences, among the 40 piRNAs they have cloned from germline of male mouse^6^. Watanabe *et al* also observed 5 motifs in the germline small RNAs (gsRNA) of mouse^5^. Our bioinformatic analysis predicted, nucleus and mitochondria as the localization signal for the first motif. Accordingly, the piRNAs are reported to be observed in mitochondria of cancer cells^32^ and nucleus of *Aplysia* neurons^18^. Out of the 102 piRNAs, 7 of them mapped to the ribosomal proteins (Table S1) implicating their probable role in rRNA processing, which is in line with the functional prediction of piRNAs. The third motif identified from our study is predicted to be involved in sensory perception which could be one of the novel functions of piRNAs in retina.

As the expression of piRNAs were reported to be tissue-specific^31^, our data also indicates that there are only 5 piRNAs in common among so far reported human tissues – brain^20^, cardiac progenitors and cardiac fibroblasts^25^, ovary^26^ and testis^27^. It is interesting to note that 75% of piRNAs in retina/RPE showed similarities with the piRNAs reported in human brain (668 piRNAs) although testis had a larger number of annotated piRNAs (11680 piRNAs) suggesting that a subset of piRNAs could be enriched in neurons. Our findings aligned with the study of Martinez *et al* which described the tissue specific expression of piRNAs with the potential of special functions in various somatic tissues^31^. Tissue enrichment analysis showed that piR-62011 is abundant in retina indicating its potential as a biomarker or a therapeutic target in retinal diseases. Although, we reported only the annotated piRNAs; identification of *de novo* piRNAs in retina and RPE should shed light on more ocular specific piRNAs. In this context, investigation of piRNAs in other non-gonadal tissues is necessary, in order to categorise the tissue specific piRNAs.

Computational annotation of piRNA target sites showed potential pleiotropic roles of piRNA in retina and RPE. It is intriguing that the majority of the identified putative targets of piRNAs are yet to be characterized for their function. The largest class of the targets is involved in signal transduction. For instance, piR-Hep1, which is conserved with piR-43771, piR-43772 and piR-60565 reduces the phosphorylation levels of Akt in hepatocellular carcinoma^50^ and Rajan *et al* speculated the role of piRNAs in Akt pathway^51^. Evidently, our initial work showed that HIWI2 silencing increased the phosphorylation of Akt and GSK3β. We have screened 43 kinases in Si-HIWI2 cells, involved in 4 different pathways – JAK/STAT, MAPK, Akt and AMPK. Apart from modulation of Akt pathway, we observed significant variation in the phosphorylation levels of STAT3, p53, AMPKα and HSP90^21^. In the similar line, Wang *et al* have also shown the influence of HIWI2 in the activation of MAPK/ERK, TGFβ and FGF signalling pathways^52^. It is plausible that these changes could be mediated through piRNAs, suggesting its involvement in cell signalling and cell-cell contact.

Other key group of identified piRNA targets were related to specific functions of retina and we have validated the expression of selective novel targets in the absence of piRNAs. Downregulation of *STX1A, Rab5, Rab8, VAMP7* and *VAMP8* indicates the likely association of piRNAs in intracellular trafficking. Rab8 is suggested to participate in ciliary trafficking of rhodopsin and in the morphogenesis of rod outer segment disk^53^. Moreover, Rab8 and Rab11 coordinates the primary ciliogenesis mainly by regulating the vesicular trafficking^54^. Primary cilia is crucial for sensory perception where defects in ciliogenesis have been observed in Bardet-Biedl syndrome (BBS) and retinal degeneration is one of the primary characteristics of BBS^55^. Rabin8 directly interacts with the BBSome^56^ which is stimulated by Rab11^54^ underscoring the importance of trafficking proteins in sensory perception. In line with our study, piR-34393 is known to target *Rab11A* in human brain^20^. Remarkably, mutations in *RPGRIP1*, an interacting protein of RPGR (retinitis pigmentosa GTPase regulator), is associated with diseases like leber congenital amaurosis, retinitis pigmentosa and cone-rod dystrophy. It is also known to be essential for the development of rod outer segments by regulating the ciliary protein trafficking and Rab8 was observed to be mislocalized in *rpgrip1* mutants^35^. Our data shows significant downregulation of both *Rab8* and *RPGRIP1* transcripts in the absence of piRNAs. Interestingly, expression of *RPGRIP1* was readily detected only in retina and testis when a panel of RNA from human tissues were screened^57^. piRNAs, which were thought to be specific to testis, might regulate *RPGRIP1* and thus may effect neurodegeneration which needs to be confirmed by further functional experiments.

LRPAP1 (LDL receptor related protein associated protein 1), another predicted piRNA target, is also downregulated in Si-HIWI2 cells. *LRPAP1*is known to be associated with high myopia by influencing the activity of TGFβ and thereby remodelling the sclera^58^. Alterations in *LRPAP1* is also related with late-onset of AD^37^. Esposito *et al* have shown the expression of piRNAs in AD^59^ and a further study demonstrated dysregulation of piRNAs in AD^20^. Together with *RPGRIP1*, downregulation of *LRPAP1* indicates a possible role of piRNAs in retinal degeneration.

Our data also showed that TIMELESS, a protein needed for the regulation of circadian rhythm in mammals, is observed to be affected in the absence of piRNAs. Activation of circadian piRNA expression is suggested to be a strategy adopted by *Drosophila melanogaster* to maintain the genomic integrity during aging^60^. Furthermore, the role of PIWIL2 in suppressing the degradation of circadian proteins CLOCK and BMAL1 through phosphorylation of Akt/GSK3β and by binding to the E-box sequences of BMAL1/CLOCK complex, confirms the importance of PIWI-like proteins in regulating circadian rhythms^61^. It is worth noting that miR-182 is known to target the 3’UTR of *CLOCK* mRNA^62,63^ and is inhibited by GSK3β^64^. Similarly, HIWI2 also affects the Akt/GSK3β pathway^21^ and as mentioned earlier, piR-62011 aligns exactly with the seed sequence of miR-182 indicating a potential functional interaction between piR-62011/miR-182 in influencing the activity of CLOCK-controlled genes. In addition, improper processing of pre-miR-182 is documented to be the major cause of depression in insomnia patients^63^. Hence, it would be interesting to explore the regulatory role played by piRNAs and miRNAs in the circadian rhythm related disorders.

It is intriguing that the examined targets were all downregulated at transcript level although they were selected based on complementarity using miRanda. It is known that piRNAs regulate the expression of genes by imperfect base-pairing rule^65^, a mechanism similar to miRNAs. The possible mechanism behind the repression of the transcripts could be epigenetic activation^66^ or silencing^18^ through repressive epigenetic marks by piRNAs at the transcriptional level. Nevertheless, we cannot rule out the mechanism of posttranscriptional regulation by piRNAs as shown in transposon-driven mRNAs^67^.

Our data showed that expression of piRNAs are elevated under oxidative stress. Martinez *et al* recently reported 40 piRNAs that were regulated by hypoxia in tumors^68^. It is possible that these piRNAs might have been influenced by the oxidative stress/reactive oxygen species generated by the hypoxic conditions. In eye, oxidative stress plays a major role in the pathogenesis of many diseases such as keratoconus, Fuchs’ endothelial corneal dystrophy, Leber’s hereditary optic neuropathy and cataractogenesis^38^. ROS and oxidative stress is also known to stimulate inflammation and pathological angiogenesis in diabetic retinopathy and it plays a pivotal role in the pathogenesis of age-related macular degeneration^38^. Thus, several evidences indicate a potential role of piRNAs in retinal pathologies.

## Conclusion

Our study showed the first line of evidence for the presence of piRNAs in human retina and RPE. We have further identified and validated novel targets of piRNAs which are essential for fundamental functions of retina and RPE. The intriguing finding of our analysis is the seed sequence similarity between piR62011 and sensory organ specific miR183/96/182 cluster. The crosstalk may modulate the unique cellular demands of retina, such as, circadian expression of phototransduction components and polarized trafficking in photoreceptors. Our study also hints the possibility of interplay between piRNAs and miRNAs in regulating gene expression at transcriptional or post transcriptional level. In addition, altered expression of piRNAs under oxidative stress implies their role in pathological conditions. In summary, our finding highlights the significance of PIWI/piRNAs in visual function and retinal degeneration.

## Materials and methods

### Data source

Small RNA sequencing datasets of human retina and RPE was downloaded from (Accession ID: E-MTAB-3792)^23^. The dataset consisted of 16 human retina and 2 RPE tissue samples. The datasets ERR973601-ERR973608 and ERR973611-ERR673618 were retinal samples. ERR973609 and ERR973610 were RPE datasets.

### Data processing

Adapter sequences present in the datasets were removed using Cutadapt^69^ with PHRED score 30 and the sequences less than 20 nt were filtered out. The reads which mapped to other non-coding RNAs (tRNA, snRNA, snoRNA, miRNA, rRNA, mtRNA) were discarded and the unmapped reads were taken for further analysis. miRNA sequences were downloaded from miRBase^70,71^ and all other RNA sequences were retrieved from Pfam^72^. The reads were mapped using BOWTIE^73^.

### Identification and differential expression of known piRNAs

After eliminating the known annotated small RNAs, unique reads and read count from the data were retrieved using unique_reads_for_mapping.pl from miRGrep tool^74^. Unique reads were aligned to the known piRNA database downloaded from piRNAquest^24^ using soap.short tool^74^ with default parameters (allowed 2bp mismatches). Output file was used to calculate the final read count using ‘awk’ command and retain the piRNAs which are having ≥10 read count in any one library. Details of the origin of piRNAs were retrieved from piRNAQuest^24^ and the piRNAs were categorised as clusters when minimum of 3 piRNAs originated within 20 Kbp.

The quantification of each piRNA in each library was normalized to TPM (Tags per million reads) and further converted to Log2 values. Before Log2 conversion, piRNAs which were having zero TPM in any one of the library were removed. Log2 values were used in Morpheus to generate heatmap.

### Tissue enrichment analysis

piRNAs annotated in brain^20^, cardiac progenitors and fibroblasts^25^, ovary^26^ and testis^27^ were downloaded. In case of human brain, piRNAs reported in both healthy and AD brain were combined and the duplicates were removed which amounted to 668 in total. For piRNAs in cardiac progenitors and cardiac fibroblasts, the piRNAs listed in cardiospheres, cardiosphere-derived cells and cardiac fibroblasts were combined and the redundant entries were eliminated resulting to 2366 piRNAs. Out of 20121 piRNAs in human testis, same piRNAs of varying lengths were considered as one entry and a total of 11680 piRNAs were compared with retina/RPE. InteractiVenn^75^ was used for tissue enrichment analysis.

### Motif prediction and lncRNA/miRNA/piRNA alignment

Motifs in the piRNA sequences were predicted using MEME software^76^ with site distribution parameter to be any number of repetitions and the putative functions of the motifs were analysed by GOMo^76^ with the default factors. Soap.short tool^74^ was used to compare the identified piRNA sequences with the lncRNA sequences downloaded from the database LNCipedia^77,78^. The alignment was performed with the default parameters allowing maximum number of 2 bp mismatches. Human miRNA sequences from miRbase were aligned with the identified piRNAs to analyse seed sequence similarity and the alignments with the maximum of 2 mismatches are listed in Supplementary table S3.

### Identification of transcription factor binding sites

We analysed the Transcription start site (TSS), promoter and TFBS for the piRNA genes. Human promoter prediction tool, FPROM^79^ was used to predict the TSS and promoter site within the 10 kb upstream of the piRNA locus, extracted from the UCSC table browser. TFBS was scanned 1kb upstream of each piRNA using P-Match 1.0^80^ with vertebrate matrix groups and the cut-off has chosen to minimize the sum of both false positive and false negative error rates.

### Prediction of piRNA targets

Targets of piRNAs were predicted using miRanda v3.3a algorithm^81–83^ with the following parameters: Gap Open Penalty (-9.0); Gap Extend Penalty (-4.0); Score Threshold (200.0); Energy Threshold (1.0 kcal/mol); Scaling Parameter (4.0). The hits which had score ≥200 and energy value ≤ −20 kcal/Mol were considered to be the putative targets.

### Ethics statement

Donor eyes were obtained from CU Shah eye bank, Sankara Nethralaya with prior approval from the Institutional Review Board and Ethics Sub-Committee of Vision Research Foundation, Chennai, India. Eye globes were collected from donors after obtaining informed consent from their family. The experiments involving usage of the ocular tissues were carried out according to the Tenets of the Declaration of Helsinki.

### Cell culture

ARPE19 cells (ATCC-CRL-2302) (ATCC, USA) and Y79 cells were cultured using DMEM-F12 (Sigma Aldrich, USA) and RPMI-1640 respectively with 10% (v/v) heat-inactivated Fetal Bovine Serum (ThermoFisher Scientific, USA) and was maintained at 37ºC in 5% CO_2_ incubator. ARPE19 cells were passaged the day before transfection whereas Y79 cells were passaged on the same day of transfection and 2 X 10^5^ cells were seeded in a 6-well plate. DsiRNA specific to 3’ UTR of HIWI2 (Integrated DNA Technologies, USA) was transfected using Lipofectamine RNAiMAX (ThermoFisher Scientific, USA) according to manufacturer’s instruction. Scrambled sequence was used as experimental control and RNA was isolated from Si-Control and Si-HIWI2 cells, 48 h post transfection. To induce oxidative stress, the cells were treated with 200 μM H_2_O_2_ for 24 h and RNA was isolated from the same to check for the expression of piRNAs.

### Real time PCR

Primers for piRNA and miRNA amplification were designed using miR2 primer software^84^. cDNA conversion was done with 1 μg of RNA based on the protocol by Busk and Cirera, 2014^85^. The cocktail for cDNA conversion contained Poly A polymerase (New England Biolabs, USA) for adding poly A tail; M-MuLV reverse transcriptase (New England Biolabs, USA) and a RT-primer of sequence - 5’-CAGGTCCAGTTTTTTTTTTTTTTTGC-3’, to convert the poly A tailed RNA to cDNA. 25 ng of cDNA was used for amplification of piRNAs listed in Table 2. Transcripts other than small RNAs were amplified from cDNA converted using iScript cDNA synthesis kit (Biorad, USA).

**Table 2.**
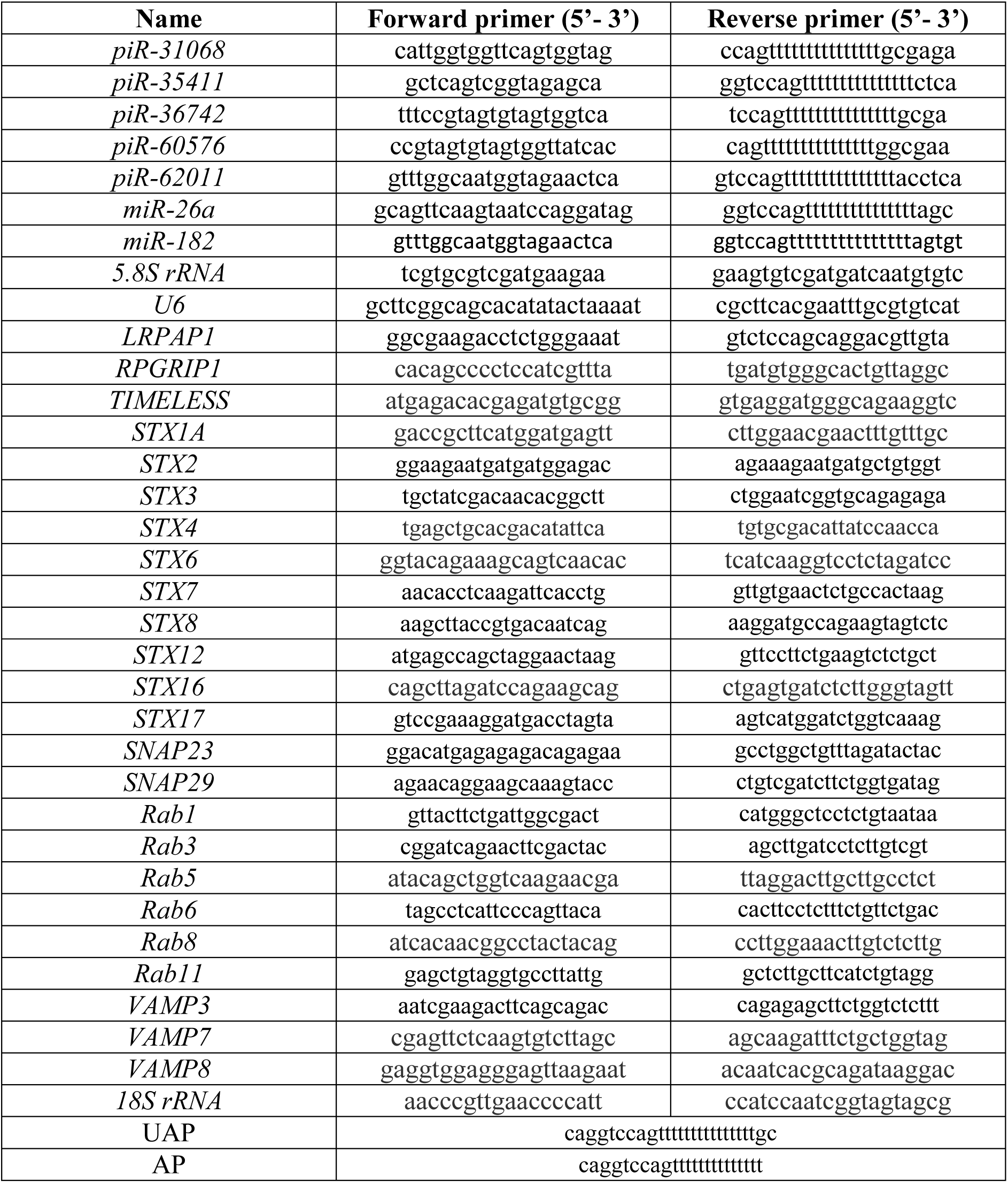
Primer sequences used for RT and qRT-PCR

### Reverse Transcription at Low dNTP concentration followed by PCR (RTL-P) analysis

RTL-P analysis was carried out as described by Dong et al, 2012^34^. RNA was converted to cDNA with low concentration of dNTP (0.4 µM) in the presence of either unanchored primer (UAP) or anchored primer (AP). The same was done at high concentration of dNTP (40 µM) as well. Four different cDNA ((1) Low dNTP with UAP; (2) Low dNTP with AP; (3) High dNTP with UAP; and (4) High dNTP with AP) were used to amplify piRNA and miRNA and the amplicons were run in 2% agarose gel. Primers used in the study are listed in Table 2.

### Western blot

After transfection, the cells were lysed using Radio Immunoprecipitation Assay (RIPA) buffer (150 mM NaCl, 0.1 %TritonX-100, 0.5 % sodium deoxycholate, 0.1%SDS, 50 mM Tris, pH 8.0) with protease inhibitors (1 mmol/L dithiothreitol, 0.5 mmol/L phenylmethylsulfonyl fluoride, 1 mg/mL leupeptin, 10 mmol/L p-nitrophenylphosphate, 10mmol/L h-glycerol phosphate) and were sonicated. The lysates were then centrifuged at 10, 000 rpm for 5 min and the proteins in the supernatant were estimated by BCA protein assay reagent (Thermo Scientific,Waltham, USA). 35 µg of protein was resolved using SDS-PAGE gel and was electrotransferred to nitrocellulose membrane (GE healthcare, UK). The blots were then incubated with 5 % blocking solution for 1 h (5 % skimmed milk powder in Tris Buffered Saline) after which they were probed for either HIWI2 (Thermo Scientific, Waltham,USA) or βACTIN (Santa Cruz Biotechnology, Dallas, USA) by incubating overnight at 4ºC with 1 in 1,000 dilution of primary antibody. Imaging of the blots were done using FluorChem C3 (Protein Simple, San Jose, USA) after incubating with the respective secondary antibodies (1 in 10, 000 dilution) for 2 h.

### Data availability

The datasets analysed were downloaded from the ArrayExpress (Accession No: E-MTAB-3792) (https://www.ebi.ac.uk/arrayexpress/experiments/E-MTAB-3792/).

## Acknowledgements

We acknowledge the financial assistance provided by Department of Science and Technology under the project “SR/FT/LS-145/2010” and Department of Biotechnology under the project DBT-Neurobiology Taskforce - “BT/PR5055/MED/30/761/2012”. The funding body is not involved in the design of the study and collection, analysis, and interpretation of data and in writing the manuscript. We thank Indian Council of Medical Research project “BMS/ FW/BIOCHEM/2015-23270/OCT-2015/24/TN/PVT” and Max Planck-India mobility grant ‘IGSTC/MPG/FS (SC)2012’. CS thanks the support from University Grants Commission, New Delhi for the award of Assistant Professorship under its Faculty Recharge Program (UGC-FRP) and for the start-up grant.

## Author contributions

SS designed, performed the *in vitro* experiments; part of *in silico* experiments and wrote the manuscript. SN performed in *silico* experiments. PG and AJ interpreted the data and reviewed the manuscript. CS conceived the idea, received the funding, planned the experiments and reviewed the manuscript. All authors read and approved the final manuscript.

## Competing interests

The authors declare no competing interests.

